# PolyAMiner-Bulk: A Machine Learning Based Bioinformatics Algorithm to Infer and Decode Alternative Polyadenylation Dynamics from bulk RNA-seq data

**DOI:** 10.1101/2023.01.23.523471

**Authors:** Venkata Soumith Jonnakuti, Eric J. Wagner, Mirjana Maletić-Savatić, Zhandong Liu, Hari Krishna Yalamanchili

## Abstract

More than half of human genes exercise alternative polyadenylation (APA) and generate mRNA transcripts with varying 3’ untranslated regions (UTR). However, current computational approaches for identifying cleavage and polyadenylation sites (C/PASs) and quantifying 3’UTR length changes from bulk RNA-seq data fail to unravel tissue- and disease-specific APA dynamics. Here, we developed a next-generation bioinformatics algorithm and application, PolyAMiner-Bulk, that utilizes an attention-based machine learning architecture and an improved vector projection-based engine to infer differential APA dynamics accurately. When applied to earlier studies, PolyAMiner-Bulk accurately identified more than twice the number of APA changes in an RBM17 knockdown bulk RNA-seq dataset compared to current generation tools. Moreover, on a separate dataset, PolyAMiner-Bulk revealed novel APA dynamics and pathways in scleroderma pathology and identified differential APA in a gene that was identified as being involved in scleroderma pathogenesis in an independent study. Lastly, we used PolyAMiner-Bulk to analyze the RNA-seq data of post-mortem prefrontal cortexes from the ROSMAP data consortium and unraveled novel APA dynamics in Alzheimer’s Disease. Our method, PolyAMiner-Bulk, creates a paradigm for future alternative polyadenylation analysis from bulk RNA-seq data.

## INTRODUCTION

Alternative polyadenylation (APA) is a post-transcriptional regulatory mechanism that cleaves a pre-mRNA molecule and appends adenosine residues at one of its potentially several cleavage and polyadenylation sites (C/PASs), ultimately resulting in multiple mRNA isoforms with varying 3’UTR lengths. By controlling the length of the 3’UTR, APA allows for the differential inclusion of binding sites specific for miRNAs and RNA-binding proteins (1). As more than half of human genes contain C/PASs and undergo APA, this widespread phenomenon plays critical roles in development and its misregulation has been implicated in several diseases, including neurodegeneration and cancer (2–4). With increasing awareness of its role in human health and disease, researchers have recognized APA as a critical post-transcriptional mechanism than previously realized. Consequently, the community has only recently developed specialized deep 3’UTR sequencing protocols such as PAC-Seq, PAS-Seq, and 3’READS to further study this phenomenon in various disease models (5–7).

However, there is an immediate need for a robust computational model that can leverage existing bulk RNA-seq datasets to decipher APA dynamics accurately and precisely. For example, multi-omic data consortiums like the Religious Orders Study/Memory and Aging Project (ROSMAP) contain robust bulk RNA-seq of the human frontal cortex for aging and Alzheimer’s Disease (8–10). However, they are notably devoid of corresponding 3’UTR sequencing datasets required for the direct study of APA dynamics. Furthermore, resequencing the >800 samples from the ROSMAP data consortia is cumbersome and impractical. Other data consortiums like The Cancer Genome Atlas (TCGA), which contains over 20,000 samples from control and primary cancer disease populations spanning 33 cancer types, and the Answer ALS data portal, which contains over 1,200 samples from control and neurodegenerative disease populations, can similarly benefit from such a tool (11, 12).

Current computational approaches for identifying C/PASs and quantifying 3’UTR length changes from bulk RNA-seq data fail to unravel tissue- and disease-specific APA dynamics (Supplementary Figure S1). The current generation of bioinformatic tools predominantly rely on (i) *a priori* C/PAS annotations, (ii) transcript reconstruction, (iii) poly(A)-capped reads, and (iv) read density fluctuations near the 3’UTR (13). Databases containing predefined *a priori* C/PAS annotations are incomplete, contain artificial noise, and do not converge with other *a priori* C/PAS databases (14–21). Methods that try to infer 3’UTR usage by transcript reconstruction from bulk RNA-seq data are hampered by inherent limitations of transcript assembly. In addition to being computationally demanding when reconstructing lowly expressed transcripts, these tools often ignore isoforms with shorter 3’UTRs as they inaccurately assign reads when shorter isoforms are embedded in longer isoforms (22, 23). Furthermore, tools that only rely on poly(A)-capped reads or reads that contain unmapped stretches of adenosines suffer from low sensitivity as these softclipped reads are relatively scarce in standard bulk RNA-seq data due to reduced read coverage near transcript ends (24). Lastly, tools whose core APA inference engine centers around detecting read density fluctuations near the 3’UTR require good coverage of the 3’UTR (25–27). This restriction limits the number of qualified genes in a sample for APA analysis after discarding genes with low read coverage. Furthermore, this class of tools is particularly vulnerable to non-biological variability and read density heterogeneity.

In sum, the current generation of bioinformatics tools for identifying C/PASs and quantifying 3’UTR length changes from bulk RNA-seq data are limited by poor C/PAS annotations that do not converge with other C/PAS databases, intrinsic limitations of *de novo* C/PAS detection, and inability to detect intra-distal or intra-proximal APA changes. Recently, an attention-based deep learning model, DNABERT, has been used to detect alternative splice sites from genomic sequences using a directional encoder representation (BERT) to capture a global understanding of genomic sequences based on neighboring nucleotide contexts (28). This landmark study showcases the power of attention-based models as they do not rely on motifs’ presence; instead, they model DNA as a language and capture hidden genomic grammar and the semantic dependency between multiple DNA sequence features. Yet, no deep learning model with a similar attention-based architecture exists to identify C/PASs. The contextual semantic insights garnered by such a model would overcome the limitations of current C/PAS databases by filtering sequence artifacts and retaining true C/PASs. Here, we develop a novel bioinformatics algorithm and application, PolyAMiner-Bulk, that addresses not only these concerns but also offers an end-to-end paradigm for the complete analysis of alternative polyadenylation changes from input bulk RNA-seq data. The methodical flow of PolyAMiner-Bulk is illustrated in Figure 1. In brief, PolyAMiner-Bulk detects *de novo* C/PASs, merges them with *a priori* C/PAS database like PolyA_DB and PolyASite, filters these candidate C/PASs using the C/PAS-BERT machine learning model to create an accurate and comprehensive C/PAS collection, and employs vector projections to examine APA dynamics throughout the gene body. A detailed description of the proposed approach is given in the following Materials and Methods section.

**Figure 1.**
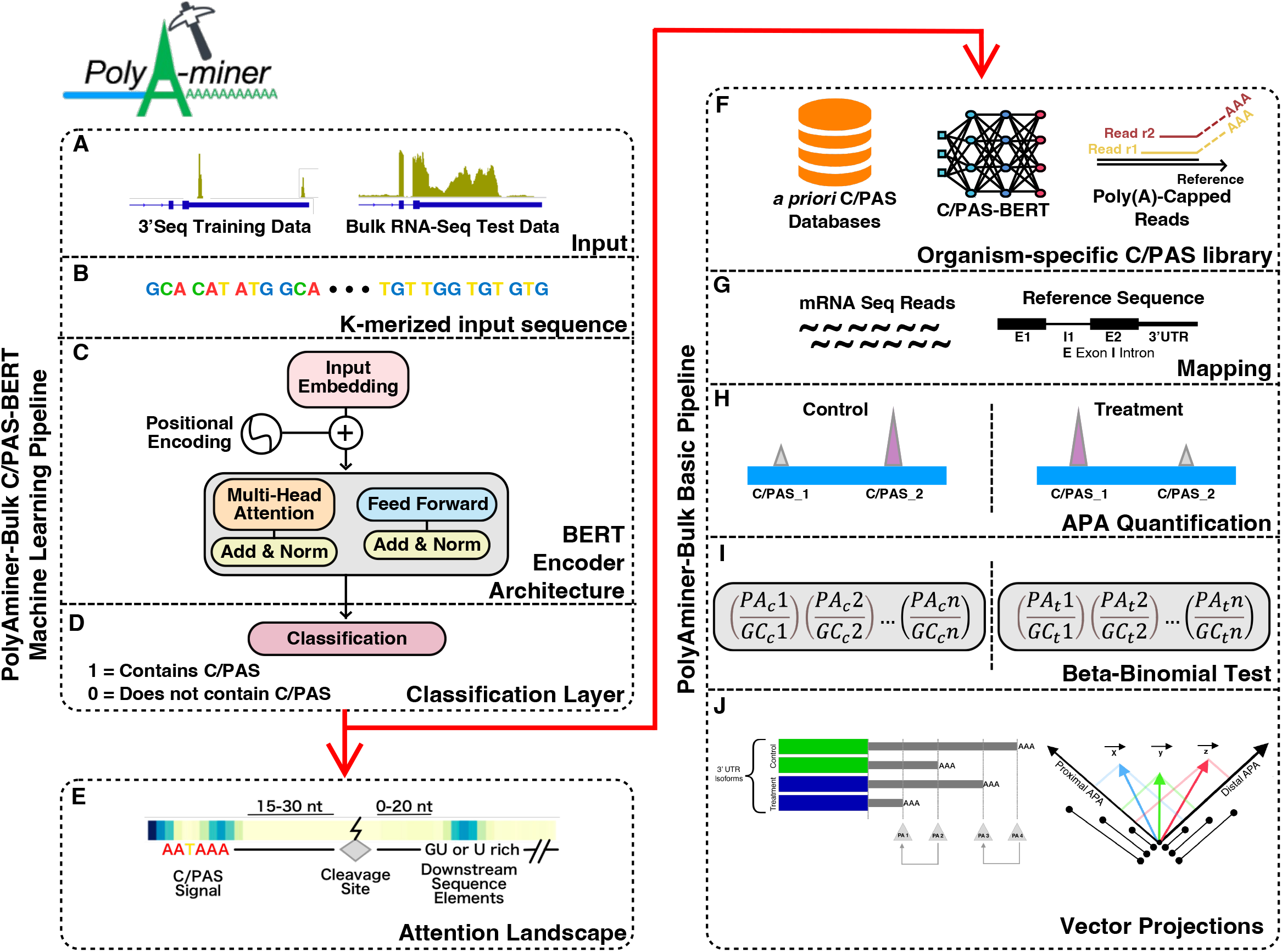
Illustration of PolyAMiner-Bulk pipeline: (A) Input data (B) K-merized input sequence (C) BERT encoder architecture (D) Classification layer (E) Attention landscape (F) Amassed and filtered candidate C/PAS library (G) Mapping (H) APA quantification (I) Beta-binomial statistical testing (J) Vector projections. In brief, PolyAMiner-Bulk detects *de novo* C/PASs, merges them with *a priori* C/PAS database like PolyA_DB and PolyASite, filters these candidate C/PASs using the C/PAS-BERT machine learning model to create an accurate and comprehensive C/PAS collection, and employs vector projections to examine APA dynamics in all genic regions.

## MATERIAL AND METHODS

### Processing raw reads

PolyAMiner-Bulk can take either the raw read files in fastq format or the mapped alignment files in bam format as input. Raw reads are mapped to the reference genome of origin using STAR and the resulting alignment files (in bam format) are sorted and indexed using samtools (29, 30).

### Extracting *de novo* C/PASs

PolyAMiner-Bulk amasses a candidate C/PAS collection from two sources: (i) directly from the input data and (ii) indirectly from existing C/PAS databases. In addition to incorporating *a priori* C/PASs into our candidate C/PAS library for downstream C/PAS-BERT mediated filtering, PolyAMiner-Bulk detects *de novo* C/PASs using softclipped read detection. The entirety of a read need not be completely aligned to a reference as the read may contain additional bases that are not in the reference or may be missing bases in the reference. This softclipped region phenomenon underscores the *de novo* C/PAS extraction engine of PolyAMiner-Bulk. Candidate *de novo* C/PASs are defined as reads from BAM read alignment files whose ends are softclipped regions containing a softclipped length-dependent proportion of adenosines (or thymines, depending on the strandedness of sequencing). For example, a softclipped tail of >12 nucleotides must contain at least 75% adenosines to be classified as a candidate *de novo* C/PAS. Shorter softclipped tails require a proportionally greater percentage of adenosines. The default settings for this user-adjustable parameter are at least 90% adenosines for a 4 nucleotides long-softclipped tail, at least 85% adenosines for a 4-8 nucleotides long-softclipped tail, at least 80% adenosines for an 8-12 nucleotides long-softclipped tail, and at least 75% adenosines for a >12 nucleotides long-softclipped tail. This loose thresholding approach is crucial as the poly(A) stretch may not necessarily continue until the end of the read because sequencing can continue into primer sequences at the end of fragments, sequencing quality of stretches of the same nucleotide may rapidly deteriorate, and sequencing errors might disrupt the poly(A) stretch.

### Clustering C/PASs

We equipped PolyAMiner-Bulk with two C/PAS clustering modes: (i) softclipped and *a priori* clustering, as well as (ii) softclipped-assisted clustering (Figure 2). *De novo* and *a priori* C/PASs are clustered in both modes based on a user-defined cluster distance parameter (default = 30 bp). Since alternative polyadenylation is a relatively non-specific process by which cleavage and polyadenylation can occur within a range of a few nucleotides from a C/PAS, PolyAMiner-Bulk selects the most distal C/PAS within a cluster. Notably, in the softclipped-assisted clustering mode, PolyAMiner-Bulk only keeps softclipped-supported clusters (Figure 2A and 2B). This mode allows for additional specificity in selecting C/PASs supported by the dataset. Other parameters are included to refine this specificity even further, such as a parameter for the minimum number of softclipped reads required for a cluster to be kept and another parameter for the minimum number of unique samples that must meet the criteria mentioned above. Figure 2C shows a representative differential APA gene identified using the softclipped-assisted clustering mode.

**Figure 2.**
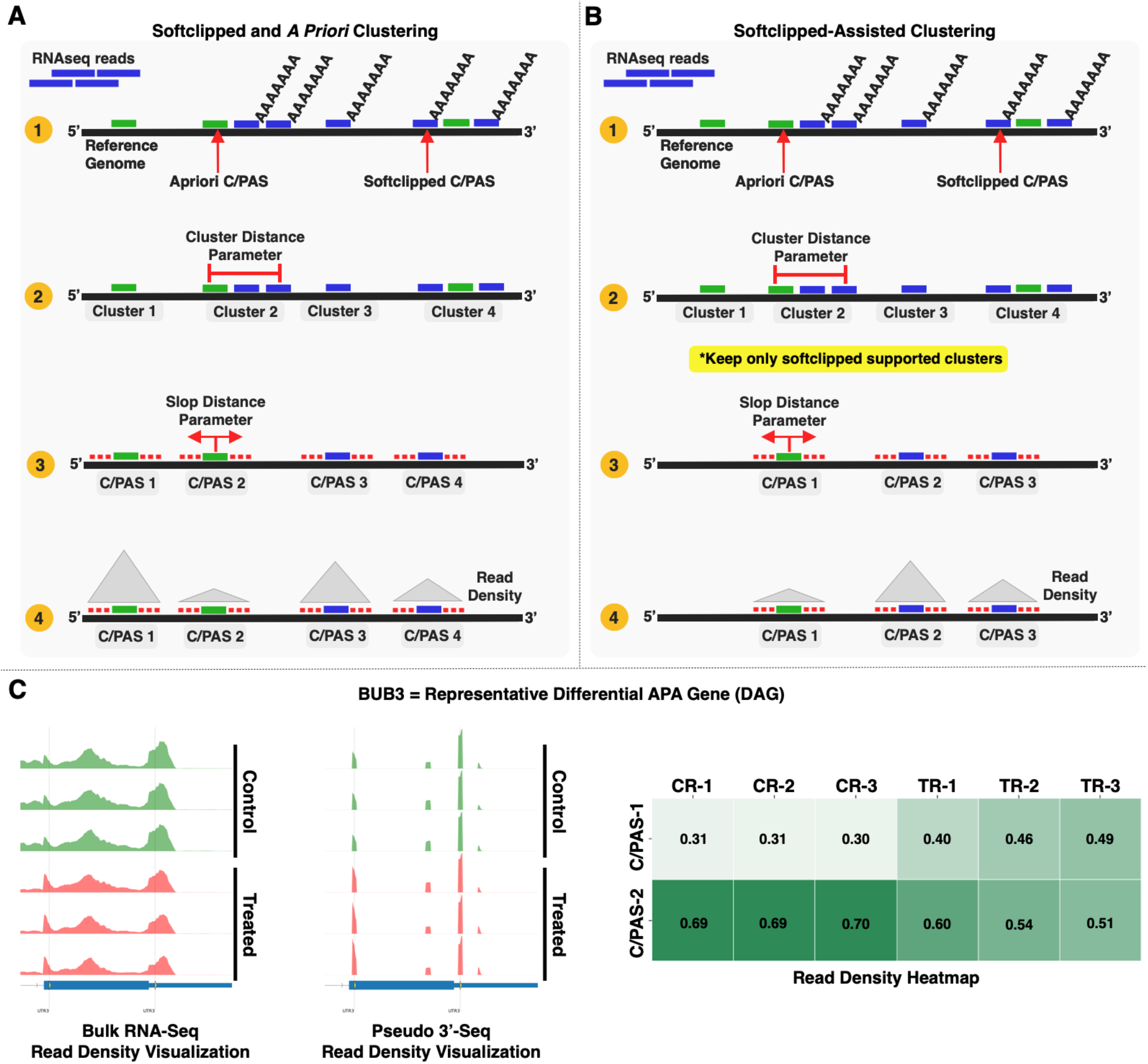
(A) Softclipped and *a priori* clustering mode. (B) Softclipped-assisted clustering mode. In this mode, PolyAMiner-Bulk only keeps softclipped-supported clusters, allowing for additional specificity in selecting C/PASs supported by the dataset. (C) Representative differential APA gene identified using the softclipped-assisted clustering mode.

### Filtering candidate C/PASs with C/PAS-BERT

When we overlapped two of the field’s most widely used predefined human-specific a priori C/PAS databases, PolyASite and PolyA_DB, we observed that both databases share 182,608 elements that constitute ∼30% of PolyASite and ∼60% of PolyA_DB (Figure 3A). This finding strongly suggests that, although both databases are based on 3’UTR-seq (rather than bulk RNA-seq) technology, they do not capture all C/PASs and most likely contain false-positive C/PAS artifacts. In addition, our in-house human brain-specific 3’UTR-seq data affirms the limitations of current *a priori* C/PAS databases (Figure 3B). Taken together, these findings showcase the limitations of current *a priori* C/PAS databases: (i) Inclusion of C/PASs that are not present in the tissue-of-interest (brain in this example) but are present in other tissues, (ii) Exclusion of novel C/PASs, and (iii) Inclusion of misprimed C/PAS artifacts.

**Figure 3.**
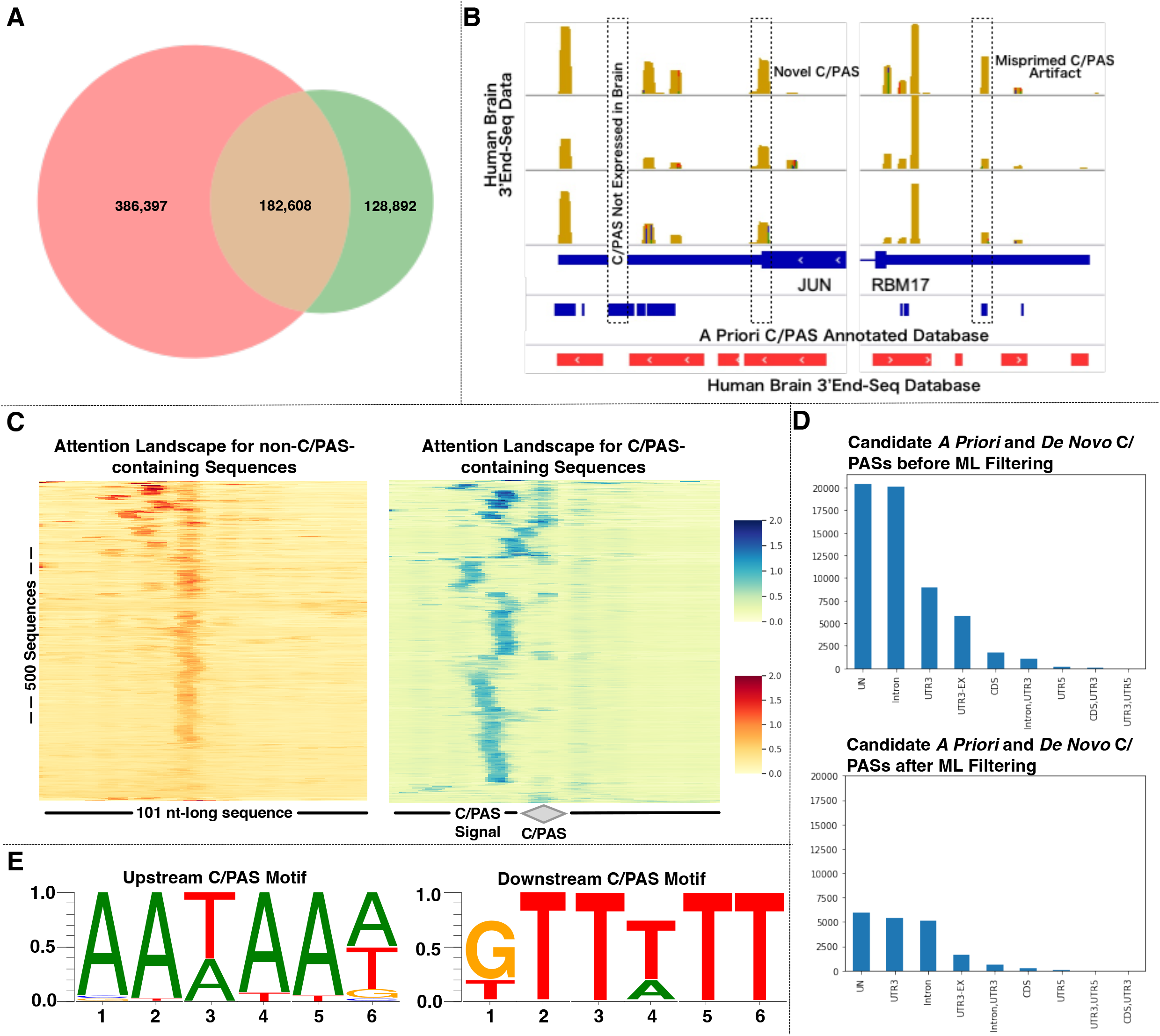
(A) Venn diagram set analysis of PolyASite and PolyA_DB and comparisons of read density visualizations of in-house human brain-specific 3’UTR-seq data with PolyASite and PolyA_DB. The limitations of current *a priori* C/PAS databases are the inclusion of C/PASs that are not present in the tissue-of-interest (brain in this example) but are present in other tissues, exclusion of novel C/PASs, and inclusion of misprimed C/PAS artifacts. (B) Overall performance metrics of the C/PAS-BERT machine learning model. (C) Attention landscapes for (non-)C/PAS-containing sequences (D) Genic C/PAS distribution before and after C/PAS-BERT filtering (E) Motif enrichment analysis on the high attention regions of C/PAS-containing sequences (F) Representative differential APA gene identified using the C/PAS-BERT model.

To filter false-positive C/PAS artifacts, we developed C/PAS-BERT, an attention-based computational model that understands distinct DNA sequence semantic relationships around C/PASs, aids in precise C/PAS identification, and improves APA inference from bulk RNA-seq data. Candidate C/PASs that both PolyASite and PolyA_DB shared were considered positively labeled C/PASs. Since the longest human 3’UTR region is less than 3000 kb, intergenic sites not within 3000 kb upstream and downstream of any annotated gene were considered negatively labeled C/PASs. 6-mer nucleotide sequence representations were first generated by querying for nucleotides that are 50 bp upstream and downstream of the candidate C/PAS and then walking over these 101 nucleotide-long DNA sequences with a 6-nucleotide long sliding window. Breaking DNA sequences into strings of every 6-nucleotide length and using them as vectors allows for sensitive and specific methods for analyzing genomes. The human dataset consisted of 633,786 tuples (6-mer nucleotide sequence representation, C/PAS label). We ensured that this dataset was balanced – the number of positively and negatively labeled tuples was equal. 90% of this overall dataset was used for k-fold cross-validation, while the remaining 10% was used as an independent test set. K-fold cross-validation was employed whereby the original dataset was equally partitioned into 12 subparts. Out of the 12 groups, for each iteration, one group was selected as validation data, and the remaining groups were selected as training data. The process was repeated until each group was treated as validation and the remaining as training data. This strategy ensures that the training process has low time complexity, the entire dataset is utilized for training and validation, and the final C/PAS-BERT model has low bias. The resultant C/PAS-BERT model helps to overcome the limitations of current C/PAS databases by filtering sequencing artifacts and better understanding APA dynamics in gene regulation.

### Statistical testing

A beta-binomial test is used to determine the significance of each PolyAIndex metric. Let J ∈ ℕ denote the number of C/PAS reads, U ∈ ℕ denote the total number of reads spanning across all C/PASs, and ℕ the set of natural numbers. Assume J is distributed according to a binomial distribution with success probability 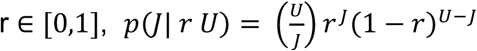. To capture the variations between biological replicates, we model r through a beta distribution with *α* > 0 and *β* > 0, *p*(*r*|*α β*) = *π*^*α*−1^(1 − *π*)^{*β*−1}^*B*(*α, β*)^−1^, where *B*(*α, β*)^−1^ is the beta function. For numerical stability, we can parametrize the beta distribution to *π* = *α*(*α* + *β*)^−1^, *ρ* = (*α* + *β*)^−1^, where *π* is the expectation of the r and *ρ* represents the dispersion. The log-likelihood of the observed data is given by:

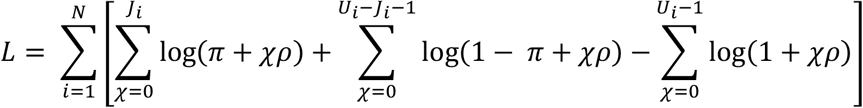

Assuming there are G groups in an experiment, we let *L*_*g*_ be the maximal log-likelihood value for group g = 1, …, G. We propose to test the homogeneity of the groups by likelihood ratio test, where the log likelihood ratio statistics S is given by 2(−*L*_0_ + ∑_*g*_ *L*_*g*_). S is approximately *χ*^2^ distribution with 2(*G* − 1) degrees of freedom. The null hypothesis of this test is that the expectation and dispersion of the different groups are equal. Every gene-level APA change is multiple testing corrected using Benjamini-Hochberg procedure (31). Gene-level APA changes with an adjusted P value < 0.05 are predicted as significant APA changes.

### Quantifying APA dynamics using vector projections

We previously deployed a vector projection-based engine to analyze differential APA dynamics from 3’sequencing data (32). We have modified this engine for PolyAMiner-Bulk to also account for the distribution of C/PASs along a gene (Figure 4A and 4B). Let us take two scenarios to illustrate the utility of this change: (i) In scenario 1, a gene shifts APA usage between two neighboring C/PASs between two conditions, and (ii) In scenario 2, a gene shifts APA usage between two faraway C/PASs between two conditions (Figure 4B). The revised engine will take the proximity of these C/PASs into account and report the gene in scenario 1 as having a smaller PolyAIndex metric, despite the gene having the same magnitude of read density change in both scenarios. This metric better reflects underlying post-transcriptional biology as a loss (or gain) of C/PASs that are farther away could result in the loss (or gain) of a more significant number of regulatory binding sites for RBPs or miRNAs. Furthermore, this vector projection-based approach we have previously pioneered accounts for ALL identified APA isoforms, unlike other methodologies that ignore APA changes involving intermediate C/PASs (Figure 4A). PolyAMiner-Bulk first projects the magnitude of C/PAS usage change to a reference C/PAS in an *n*-dimensional vector space, where *n* is the number of C/PASs. Then, it computes the difference in projections of these vectors and collapses them into a single gene-level magnitude PolyAIndex metric (Figure 4C). A positive PolyAIndex metric suggests overall 3’UTR lengthening, while a negative PolyAIndex metric suggests overall 3’UTR shortening.

**Figure 4.**
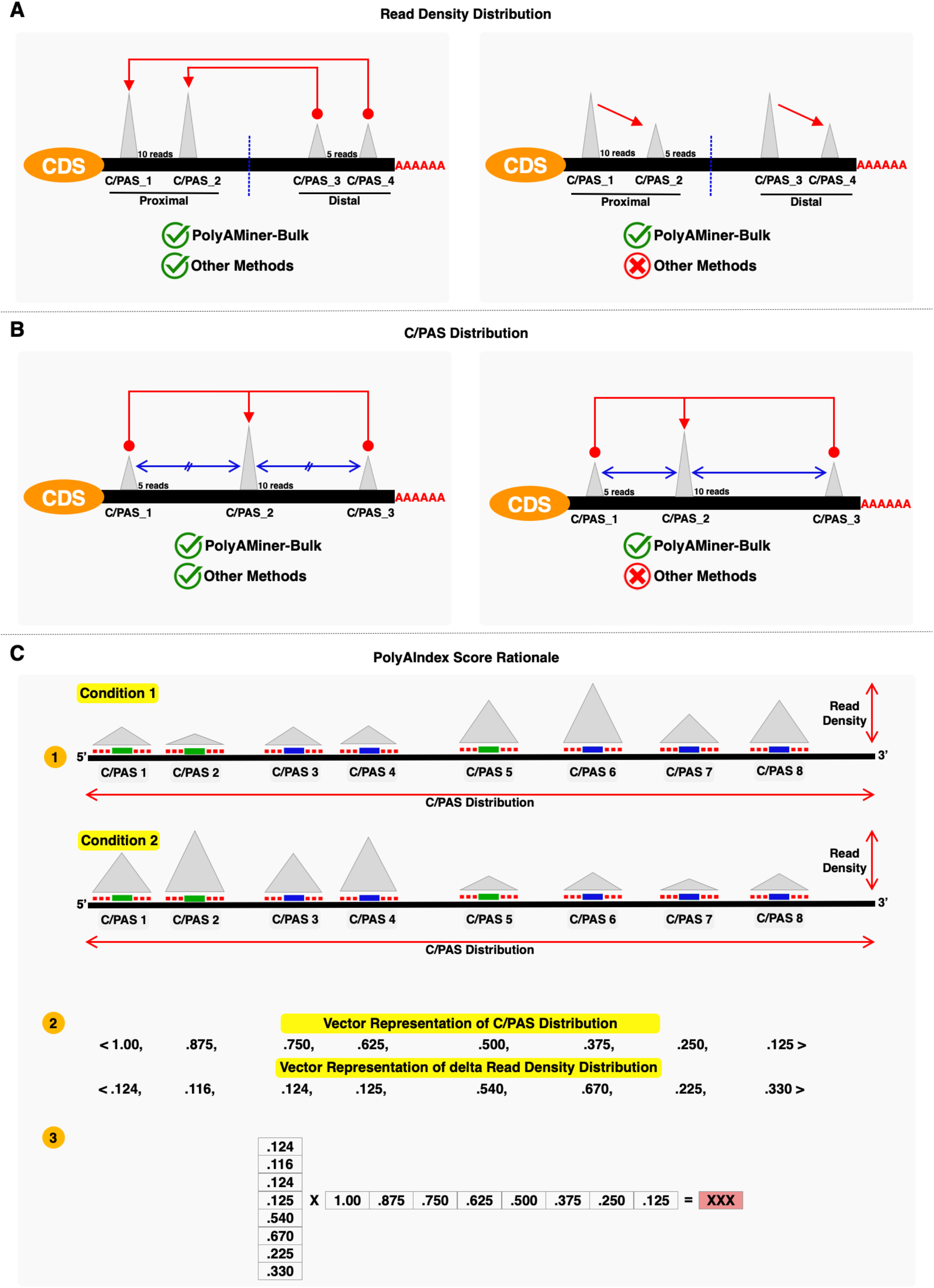
(A) Taking read density distribution accounts for ALL identified APA isoforms, unlike every other methodology that ignores APA changes involving intermediate C/PASs. (B) Taking C/PAS distribution accounts for non-equidistant 3’UTR length changes, unlike every other methodology that assumes C/PASs to be uniformly distributed. (C) Modified vector projection-based engine that takes both the distribution of C/PASs along a gene and the read density underlaying each C/PAS into account.

### Visualizing APA changes

After calculating PolyAIndex metrics, PolyAMiner-Bulk can further investigate APA dynamics of individual genes through its visualization module. We implement pyGenomeTracks and Matplotlib to generate gene-level read density coverage plots and corresponding C/PAS usage heatmaps from the bulk RNA-seq input data (33–35). Notably, we generate two gene-level read density coverage plots: (i) one showing the entire bulk RNA-seq read density coverage and (ii) the other showing the C/PAS subset of the read density to mimic 3’UTR read density coverage.

## RESULTS AND DISCUSSION

### The attention-based C/PAS-BERT machine learning model successfully identifies C/PASs and recapitulates the underlying C/PAS grammar

The overall performance was as follows: Accuracy = 0.904, Area under the curve = 0.960, F1 = 0.904, Precision = 0.904, and Recall = 0.904. Further, we visualized important regions from positively labeled (C/PAS-containing)- and negatively labeled (non-C/PAS-containing)-sequences as an attention landscape to enhance the interpretability of the C/PAS-BERT machine learning model. To visualize the attention landscape for each input sequence, we scored each nucleotide of the input sequence using the self-attention mechanism. First, we extracted the attention of the “entire sequence” on the k-mer subsequences and used it as an importance measure. Then, we converted the attention score from k-mer to the individual nucleotide by averaging the attention scores for all k-mers that contain the nucleotide. Lastly, we plotted attention for individual nucleotides as a heatmap for direct visualization (Figure 3C). Using this methodology, we retrieved two attention landscapes – one for 101-bp non-C/PAS-containing sequences and another for 101-bp C/PAS-containing sequences (where the C/PAS was located directly at the center of the sequence). Based on the non-C/PAS-containing sequence attention landscape, we observed that the model placed consistently high attention at the center of the sequence. Based on the C/PAS-containing sequence attention landscape, we can appreciate that the model learned three important genomic features previously validated as necessary for C/PAS detection. We observed consistently high attention upon (i) the C/PAS, which is located at the center of the sequence, denoted henceforth as position 0, (ii) the 15-30 nucleotide region upstream from the C/PAS containing C/PAS signal motifs like AATAA, and (iii) GT- or T-rich sequence elements that are located 0-20 nucleotides downstream from the C/PAS. This data demonstrates the power of attention-based deep learning models as they do not rely on motifs’ simple presence or absence. Rather, they employ powerful contextual understandings to understand multiple semantic features simultaneously.

We also plotted the distribution of C/PAS locations before and after filtering (Figure 3D). Before filtering, the majority of C/PASs were in unannotated gene regions (∼20,000), intronic regions (∼20,000), and UTR3 (∼8,000). After filtering, the majority of C/PASs were in unannotated gene regions (∼6,000), intronic regions (∼5,000), and UTR3 (∼5,000). This data demonstrates that most filtered C/PASs were from unannotated gene and intronic regions. This finding aligns with expectations as most artificial C/PASs would be found in those regions while most C/PASs in the 3’UTR would be preserved.

Additionally, we performed motif enrichment analysis on the 15-30 nucleotide region upstream from the C/PAS to determine the active motifs of the C/PAS-BERT model (Figure 3E, left). We identified AATAAA and its variants (e.g., AAATAA, ATAAAA, ATTAAA, ATAAAT, ATAAAG, CAATAA, TAATAA, ATAAAC, AAAATA, AAAAAA, AAAAAT) as the upstream poly(A) signal. Moreover, a similar analysis on the nucleotide region downstream from the C/PAS yielded GTTTTT and its variants as the downstream poly(A) signal (Figure 3E, right). Retrieving these well-conserved signals increases our confidence that C/PAS-BERT is learning the necessary features to identify C/PASs. Supplementary Figure S2 shows a representative differential APA gene identified using C/PAS-BERT. Taken together, this attention-based computational model understands distinct DNA sequence semantic relationships around C/PASs and can aid in precise C/PAS identification.

### PolyAMiner-Bulk significantly enhances our ability to decode APA dynamics from bulk RNA-sequencing data

We benchmarked PolyAMiner-Bulk against several of the most common bulk RNA-seq-based APA methods to examine the APA dynamics in a bulk RNA-seq dataset of immortalized human embryonic kidney (HEK293) cells with and without siRNA-mediated knockdown of RNA-binding motif protein 17 (RBM17) (GEO: GSE107648). Previous research has shown this protein to regulate the expression and splicing of RNA-processing proteins (36). Moreover, this RBM17 knockdown and control contrast reveals differential expression of several protein factors that facilitate alternative polyadenylation (37). These differential core APA factors include NUDT21, CSTF3, FIP1L1, PPP1CB, CPSF3, PPP1CA, PAPOLA, CSTF2, PABPN1, RBBP6, and PCF11 (Figure 5A). Since these factors aid in the regulation, detection, cleavage, and polyadenylation of a C/PAS, their differential expression strongly suggests that the knockdown of RBM17 substantially perturbs APA dynamics.

**Figure 5.**
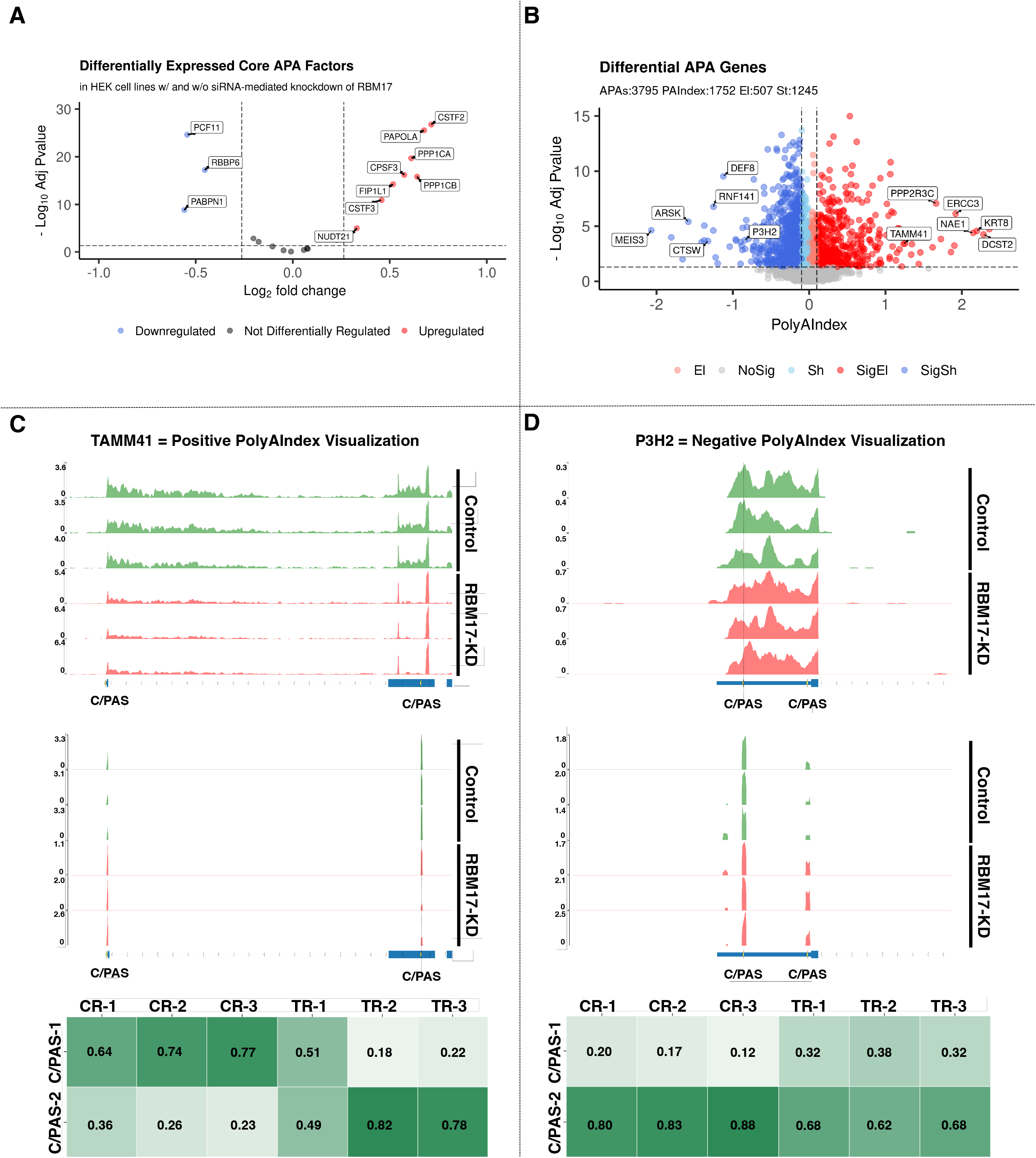
PolyAMiner-Bulk analysis of bulk RNA-seq benchmarking dataset of HEK293 cells with and without siRNA-mediated knockdown of RBM17. (A) Volcano plot of differentially expressed core APA factors. (B) Volcano plot of differential APA genes. (C) Representative differential APA gene with a positive PolyAIndex, suggesting 3’UTR elongation. (D) Representative differential APA gene with a negative PolyAIndex, suggesting 3’UTR shortening.

PolyAMiner-Bulk detected 3,795 significant differential APA genes (DAGs), of which 1752 genes exhibited a PolyAIndex magnitude greater than 0.1 or less than -0.1. Of these DAGs, 1245 underwent 3’UTR shortening, and 507 underwent 3’UTR lengthening (Figure 5B). These results are well represented by visualizations of read density fluctuations of representative DAGs near their respective C/PASs for control and RBM17 knockdown groups (Figure 5C and 5D). For example, TAMM41, involved in mitochondrial translocator assembly and maintenance, is a representative differential APA gene with a positive PolyAIndex metric, suggesting that this gene is undergoing 3’UTR lengthening in the RBM17 knockdown condition compared to the control condition (Figure 5C) (38, 39). On the other hand, P3H2, which is involved in collagen chain assembly and stability, is a representative differential APA gene with a negative PolyAIndex metric, suggesting that this gene is undergoing 3’UTR shortening in the RBM17 knockdown condition compared to the control condition (Figure 5D) (40, 41). We visualized both genes’ read density – as bulk RNA-seq and pseudo-3’UTR-seq read coverage – and plotted their corresponding density proportions as a heatmap. In the control condition, TAMM41 exhibits higher read proportion density in its proximal 3’UTR C/PAS, whereas TAMM41 shifts a proportion of its read density towards the distal 3’UTR C/PAS in the RBM17 knockdown condition. By contrast, in the control condition, P3H2 exhibits higher read proportion density in its distal 3’UTR C/PAS, whereas P3H2 shifts a proportion of its read density towards the proximal 3’UTR C/PAS in the RBM17 knockdown condition.

To assess the performance of PolyAMiner-Bulk, we tested DaPars, APAlyzer, and TAPAS, the currently utilized bulk RNA-seq-based APA methods, on this RBM17 knockdown bulk RNA-seq dataset (19, 25, 27). DaPars identified 155 differential APA genes (12 undergoing 3’UTR lengthening and 143 undergoing 3’UTR shortening); APAlyzer identified 157 differential APA genes (30 undergoing 3’UTR lengthening and 127 undergoing 3’UTR shortening); and TAPAS identified 546 differential APA genes (205 undergoing 3’UTR lengthening and 341 undergoing 3’UTR shortening). After performing an Upset Plot set analysis, we observed that PolyAMiner-Bulk not only identifies the largest number of unique differential APA genes but also identifies the highest number of differential APA genes that were also identified by other methods (Figure 6A). This increased sensitivity can be attributed to (i) our tool’s ability to detect a more accurate C/PASs and (ii) our vector projection-based approach that helps identify significant intra-distal and intra-proximal APA changes that current generation methods would have otherwise ignored. Capturing and quantifying these APA dynamics may be biologically relevant as a loss (or gain) of these intra-distal and intra-proximal C/PASs can lead to a loss (or gain) of a more significant number of regulatory binding sites for regulatory molecules like RBPs or miRNAs.

**Figure 6.**
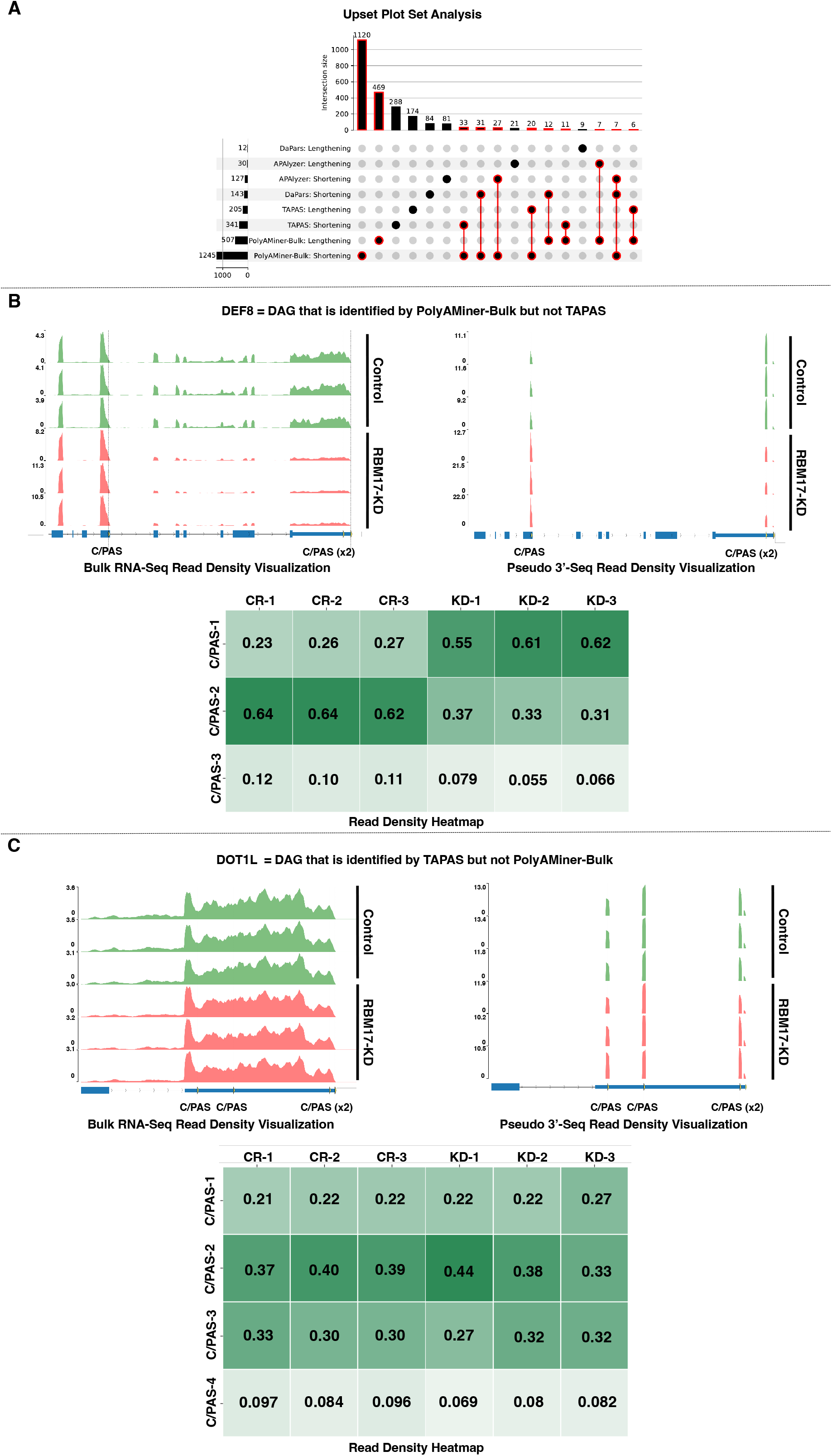
Comparison of PolyAMiner-Bulk against current-generation tools. (A) Upset plot analysis. (B) Representative differential APA gene identified by PolyAMiner-Bulk but not TAPAS, a current-generation tool. (C) Representative differential APA gene identified by TAPAS and not PolyAMiner-Bulk.

Furthermore, the differential APA genes uniquely identified by PolyAMiner-Bulk are well represented by visualizations of read density fluctuations of genes near their respective C/PASs for control and RBM17 knockdown groups. DEF8, involved in cation binding, is a representative gene classified by PolyAMiner-Bulk but not observed in other methods like TAPAS (Figure 6B) (42). Compared to the control condition, DEF8 undergoes 3’UTR shortening in the RBM17 knockdown condition. In the control condition, DEF8 exhibits higher read proportion density in its distal 3’UTR C/PAS, whereas DEF8 shifts a proportion of its read density towards the proximal 3’UTR C/PAS in the RBM17 knockdown condition. We also visualized genes uniquely identified as undergoing significant APA changes by methods other than PolyAMiner-Bulk like TAPAS. DOT1L, involved in methylating lysine-79 of histone H3 in nucleosomes, is one such representative gene (Figure 6C) (43). PolyAMiner-Bulk does not classify DOT1L as a differential APA gene since the three samples do not uniformly undergo changes in read density among the four C/PASs between each condition. The corresponding read density and heatmap visualizations further corroborate this result. Comparisons between PolyAMiner-Bulk and other methods like APAlyzer and DaPars demonstrate a similar pattern where read density visualizations better support the results of PolyAMiner-Bulk (Supplementary Figure S3).

### Revisiting published data using PolyAMiner-Bulk reveals novel APA dynamics and pathways in scleroderma pathology

Previously published studies have established that NUDT21 (also known as CFlm25), a core APA factor, directs differential alternative polyadenylation and that its suppression induces a collection of 3’UTR shortening events through loss of stimulation of distal C/PASs (44–48). One such study examined the effects of NUDT21 knockdown in normal skin fibroblasts and noted the 3’UTR shortening of key TGF-beta-regulated fibrotic genes (49). We used PolyAMiner-Bulk to re-analyze this bulk RNA-seq dataset of skin fibroblasts with and without siRNA-mediated knockdown of NUDT21 (GEO: GSE137276) and compared the output with previously published results.

PolyAMiner-Bulk detected 3,731 significant differential APA genes (DAGs), of which 2154 genes exhibited a PolyAIndex magnitude greater than 0.1. Of these DAGs, 1791 underwent 3’UTR shortening, and 363 underwent 3’UTR lengthening (Figure 7A). In contrast, the previously published study reported only 1038 DAGs, with 947 undergoing 3’UTR shortening and 91 undergoing 3’UTR lengthening (Figure 7B). Of note, PolyAMiner-Bulk not only recapitulated >50% of the 3’UTR shortening DAGs identified by the other study but also identified 1287 unique 3’UTR shortening DAGs. As discussed previously, our improved C/PAS identification paradigm and vector projection-based approach underlie PolyAMiner-Bulk’s increased sensitivity.

**Figure 7.**
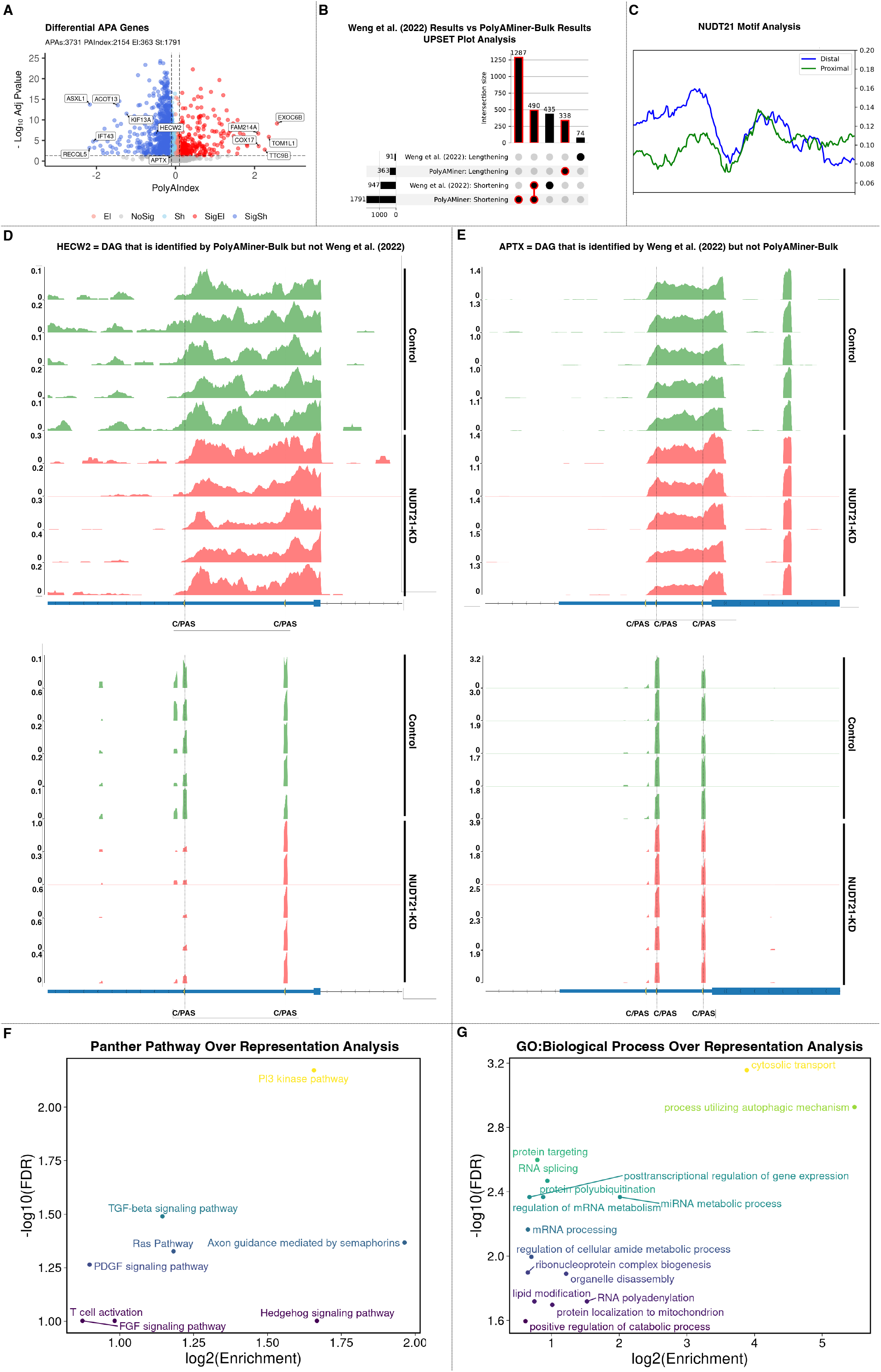
PolyAMiner-Bulk analysis of bulk RNA-seq dataset of skin fibroblasts with and without siRNA-mediated knockdown of NUDT21. (A) Volcano plot of differential APA genes. (B) Upset plot analysis of PolyAMiner-Bulk results against previously published results. (C) NUDT21 motif and biding analysis on the unique 3’UTR shortening genes identified by PolyAMiner-Bulk. (D) Representative differential APA gene identified by PolyAMiner-Bulk but not the previously published study. (E) Representative differential APA gene identified by the previously published study but not PolyAMiner-Bulk. (F) Panther Pathway over representation analysis of the differential APA genes identified by PolyAMiner-Bulk. (G) GO:Biological Process over representation analysis of the differential APA genes identified by PolyAMiner-Bulk.

To better understand the genes uniquely identified as undergoing 3’UTR shortening by PolyAMiner-Bulk, we examined the distribution of the NUDT21 binding motif, UGUA, within the 3’UTR of mRNAs (Figure 7C). Also, we compared our results with published cross-linked immunoprecipitation sequencing (CLIP-seq) data for NUDT21 in human HEK293T cells (GEO: GSE37398) (Supplementary Figure S4) (50). We found significant enrichment for UGUA motifs near distal C/PASs compared to the proximal C/PASs within the 3’UTR of genes that exhibit 3’UTR shortening changes following NUDT21 knockdown. Furthermore, after comparing our results with published CLIP-seq data for NUDT21 in human HEK293T cells, our results indicate that NUDT21 show higher and more concentrated binding at distal C/PASs of the unique 3’UTR shortening genes identified by PolyAMiner-Bulk. This binding pattern is consistent with the observed distribution pattern of the UGUA motif and supports a model whereby NUDT21 is directed to distal sites to facilitate alternative polyadenylation. Taken together, these concordant findings suggest that the 1287 unique 3’UTR shortening DAGs identified by PolyAMiner-Bulk are indeed targets of NUDT21 and actual signals.

Furthermore, PolyAMiner-Bulk results are well represented by visualizations of read density fluctuations of representative DAGs near their respective C/PASs for control and NUDT21 knockdown conditions. For example, HECW2, involved in ubiquitin-protein ligase activity, is a representative DAG uniquely identified by PolyAMiner-Bulk (51–53). This gene underwent 3’UTR shortening in the NUDT21 knockdown condition compared to the control condition – a finding that is corroborated by read density visualizations (Figure 7D). In the control condition, HECW2 exhibits higher read proportion density in its distal 3’UTR C/PAS, whereas HECW2 shifts a proportion of its read density towards the proximal 3’UTR C/PAS in the NUDT21 knockdown condition. Of significant interest, the authors of this previously published study themselves have independently identified HECW2 as being involved in scleroderma pathogenesis in a separate study (54). We also visualized genes uniquely identified by the previously published study. APTX, involved in single-stranded DNA repair, is one such representative DAG (55). PolyAMiner-Bulk does not classify APTX as a differential APA gene since the five samples for each condition do not uniformly undergo changes in read density among the three C/PASs. The corresponding read density and heatmap visualizations further corroborate this result (Figure 7E).

Lastly, we performed functional enrichment analyses to compare the biological insights from the DAGs identified by PolyAMiner-Bulk versus those identified by the previously published study. We first determined if any subset within the 3’UTR shortening DAGs identified by PolyAMiner-Bulk shared more or fewer genes with the “Panther Pathway” database than one would expect by chance. While several pathways like TGB-beta signaling and T-cell activation were enriched by both PolyAMiner-Bulk and the previously published study, other pathways like PI3 kinase, Ras, Hedgehog, FGF, and PDGF signaling pathways were uniquely enriched in the PolyAMiner-Bulk 3’UTR shortening DAG set (Figure 7F). Furthermore, over-representation functional analysis against the “Gene Ontology: Biological Processes” database reveals significant enrichment for posttranscriptional regulation of gene expression, protein polyubiquitination, mRNA processing, and positive regulation of catabolic process in the PolyAMiner-Bulk 3’UTR shortening DAG set (Figure 7G). Taken together, these results demonstrate that identifying these additional DAGs increases our understanding of the underlying biology and reveals previously underappreciated APA dynamics in scleroderma.

### PolyAMiner-Bulk unravels novel APA dynamics in Alzheimer’s Disease

To demonstrate the applicability and scientific potential of PolyAMiner-Bulk to large scale data consortia, we used PolyAMiner-Bulk to explore the extent of differential APA dynamics in the post-mortem prefrontal cortexes of four control and four Alzheimer’s Disease (AD) patients from the ROSMAP data consortium. Studies that have comprehensively profiled AD brain tissue revealed a strikingly weak correlation between mRNA and protein levels among various AD models (including human postmortem brain tissue) (56, 57). For example, in our recent study of the aging brain transcriptome and proteome in a *Drosophila* model of tauopathy, we observed 42% of Tau-induced mRNA transcripts showing the opposite direction of proteome change (56). These discordant findings likely reflect feedback interactions like post-transcriptional regulation that maintain protein homeostasis, consistent with findings that transcript-protein concordance varies among coregulated gene sets. In another study, we observed 3’UTR shortening of mRNA isoforms strongly correlating with increased levels of corresponding proteins in murine neurodegenerative models (58). This finding suggests that APA may partly explain this transcriptome-proteome discordance, implicating AD as a prime disease-of-interest to investigate further with PolyAMiner-Bulk. Of note, this AD and control contrast also reveals differential expression of several core APA factors, including CPSF3, WDR33, PPP1CB, RBBP6, CSTF3, and CPSF2 (Figure 8A). Since these factors aid in the regulation, detection, cleavage, and polyadenylation of a C/PAS, their differential expression strongly suggests substantially perturbed APA dynamics.

**Figure 8.**
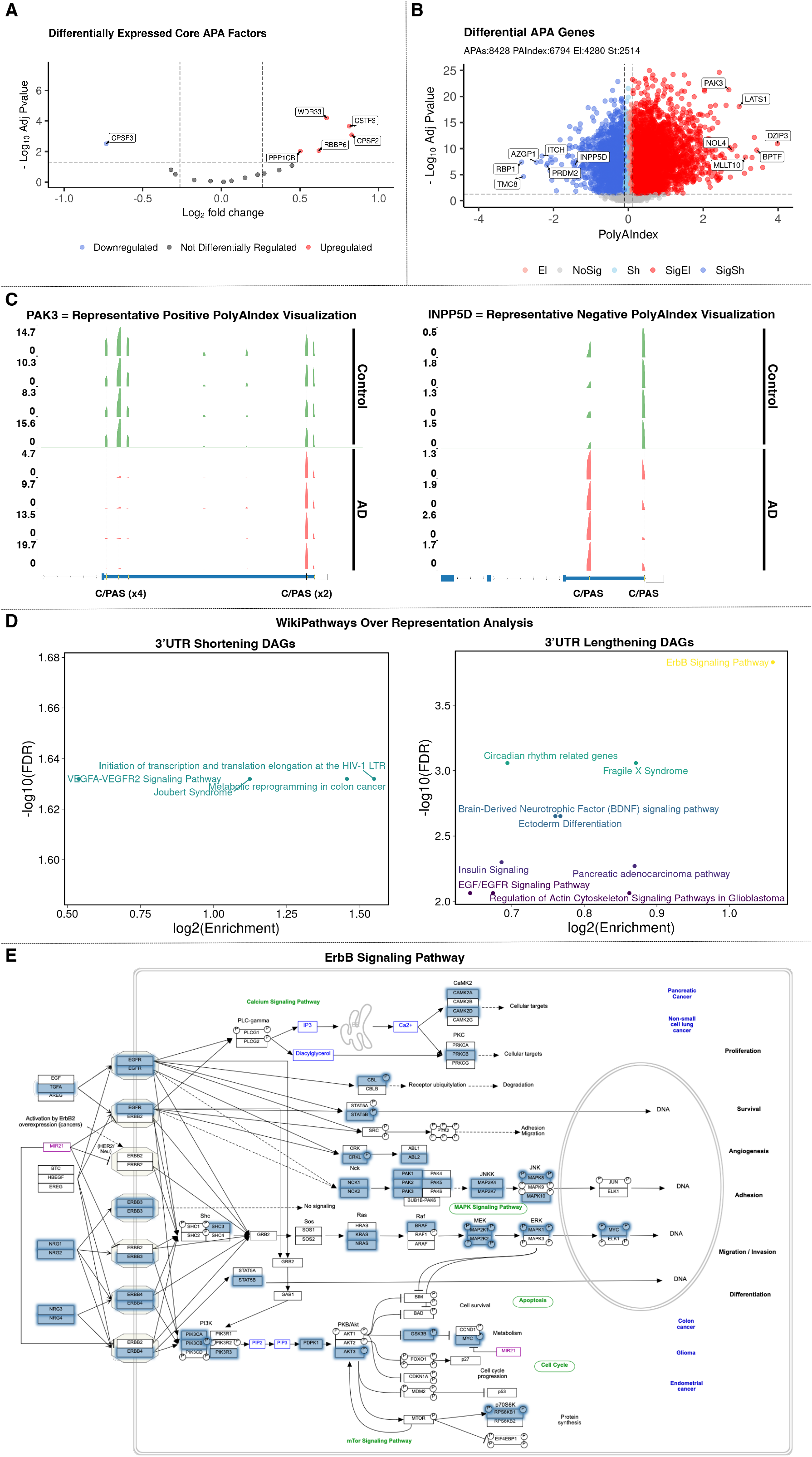
PolyAMiner-Bulk analysis of bulk RNA-seq dataset of post-mortem prefrontal cortexes of four control and four AD patients from the ROSMAP data consortium. (A) Volcano plot of differentially expressed core APA factors. (B) Volcano plot of differential APA genes. (C, left) Representative differential APA gene with a positive PolyAIndex, suggesting 3’UTR elongation. (C, right) Representative differential APA gene with a negative PolyAIndex, suggesting 3’UTR shortening. (D) WikiPathways over representation analysis of 3’UTR shortening and lengthening differential APA genes. (E) Representative enriched signaling pathway.

Aligning with our expectations, PolyAMiner-Bulk detected 8,428 significant differential APA genes (DAGs), of which 6,794 genes exhibited a PolyAIndex magnitude greater than 0.1. Of these DAGs, 2514 underwent 3’UTR shortening, and 4280 underwent 3’UTR lengthening (Figure 8B). These results are well represented by visualizations of read density fluctuations of representative DAGs near their respective C/PASs for control and AD groups. For example, PAK3, involved in dendritic development and for the rapid cytoskeletal reorganization in dendritic spines associated with synaptic plasticity, is a representative differential APA gene with a positive PolyAIndex metric, suggesting that this gene is undergoing 3’UTR lengthening in the AD condition compared to the control condition (Figure 8C, left panel) (59–61). On the other hand, INPP5D, which is involved in the negative regulation of myeloid cell proliferation and survival, is a representative differential APA gene with a negative PolyAIndex metric, suggesting that this gene is undergoing 3’UTR shortening in the AD condition compared to the control condition (Figure 8C, right panel) (62–66). We visualized both genes’ read density as pseudo-3’UTR-seq read coverage. In the control condition, PAK3 exhibits higher read proportion density in its proximal 3’UTR C/PAS, whereas PAK3 shifts a proportion of its read density towards the distal 3’UTR C/PAS in the AD condition. By contrast, in the control condition, INPP5D exhibits higher read proportion density in its distal 3’UTR C/PAS, whereas INPP5D shifts a proportion of its read density towards the proximal 3’UTR C/PAS in the AD condition. Also of note, previous literature has described the involvement of both genes in AD pathogenesis.

Apart from identifying interesting individual genes like PAK3 and INPP5D, we sought to unravel prevalent biological themes in the DAGs undergoing 3’UTR shortening and those undergoing 3’UTR lengthening. Therefore, we performed an over-representation functional analysis to determine if any genes from each set shared more or fewer genes with the “Panther Pathway” database than one would expect by chance. The 3’UTR shortening DAG set was significantly enriched for the VEGFA-VEGFR2 signaling pathway. In contrast, the 3’UTR lengthening DAG set was significantly enriched for ErbB, brain-derived neurotrophic factor, circadian rhythm, EGF/EGFR, and insulin signaling pathways (Figure 8D and 8E). Of note, studies have already linked these pathways to AD pathogenesis, but none have looked at these pathways from an APA-centric perspective (67–76). These results lay the preliminary groundwork for future studies to dissect the mechanism by which dysregulated APA drives AD pathobiology.

## CONCLUSION

PolyAMiner-Bulk is distinct from current-generation tools in many aspects (summarized in Supplementary Table S1). For instance, to our knowledge, this is the first tool with an attention-based machine learning architecture to identify C/PASs. Attention-based models do not rely on motifs’ presence; instead, they model DNA as a language and capture the hidden grammar and the semantic dependency between multiple DNA sequence features. The contextual semantic insights garnered by such a model overcome the limitations of current C/PAS databases by separating sequencing artifacts and other noise from the true C/PASs. Furthermore, using vector projections, PolyAMiner-Bulk accounts for all APA changes, including non-proximal to non-distal changes, and can distinguish the most distal to most proximal changes from most distal to intermediate site changes irrespective of absolute change magnitude. This sensitivity is crucial to estimating the true breadth of 3’UTR shortening and elongation. In addition, our tool takes raw FASTQ or processed alignment files as input and offers an end-to-end APA analysis paradigm. PolyAMiner-Bulk not only identifies differential APA genes but also generates (i) read proportion heatmaps and (ii) read density visualizations of the corresponding bulk RNA-seq tracks and pseudo-3’UTR-seq tracks, allowing users to appreciate the differential APA dynamics.

Analysis of bulk RNA-seq datasets of HEK cells with and without siRNA-mediated knockdown of RBM17, skin fibroblasts with and without siRNA-mediated knockdown of NUDT21, and post-mortem human prefrontal cortexes with and without AD strongly supports the value of PolyAMiner-Bulk as we demonstrated a substantial increase in the number of dynamic APA events detected. With the emerging importance of APA in understanding development and disease, PolyAMiner-Bulk can significantly improve APA analysis from bulk RNA-seq data and lead to a better understanding of APA dynamics across various diseases, from systemic sclerosis to neurodegeneration.

## DATA AVAILABILITY

PolyAMiner-Bulk is an open-source algorithm that has been packaged as an end-to-end Python application. It is freely available at https://github.com/venkatajonnakuti/PolyAMiner-Bulk.The data underlying this article are available in the Gene Expression Omnibus Database at https://www.ncbi.nlm.nih.gov/geo and can be accessed with GEO: GSE107648, GSE137276, and GSE37398. The other data underlying this article are available in the article and its online supplementary material.

## ACKNOWLEDGEMENT

We are thankful to our colleagues at the Baylor College of Medicine, Texas Children’s Hospital, and the Jan and Dan Duncan Neurological Research Institute who provided expertise that greatly assisted this research.

## REFERENCES

1. Mitschka, S. and Mayr, C. (2022) Context-specific regulation and function of mRNA alternative polyadenylation. Nat Rev Mol Cell Biol, 10.1038/S41580-022-00507-5.

2. Yuan, F., Hankey, W., Wagner, E.J., Li, W. and Wang, Q. (2021) Alternative polyadenylation of mRNA and its role in cancer. Genes Dis, 8, 61–72.

3. Patel, R., Brophy, C., Hickling, M., Neve, J. and Furger, A. (2019) Alternative cleavage and polyadenylation of genes associated with protein turnover and mitochondrial function are deregulated in Parkinson’s, Alzheimer’s and ALS disease. BMC Medical Genomics 2019 12:1, 12, 1–14.

4. Agarwal, V., Lopez-Darwin, S., Kelley, D.R. and Shendure, J. (2021) The landscape of alternative polyadenylation in single cells of the developing mouse embryo. Nature Communications 2021 12:1, 12, 1–12.

5. Routh, A., Ji, P., Jaworski, E., Xia, Z., Li, W. and Wagner, E.J. (2017) Poly(A)-ClickSeq: Clickchemistry for next-generation 3’-end sequencing without RNA enrichment or fragmentation. Nucleic Acids Res, 45, 1–16.

6. Hoque, M., Ji, Z., Zheng, D., Luo, W., Li, W., You, B., Park, J.Y., Yehia, G. and Tian, B. (2013) Analysis of alternative cleavage and polyadenylation by 3’ region extraction and deep sequencing. Nat Methods, 10, 133–139.

7. Shepard, P.J., Choi, E.A., Lu, J., Flanagan, L.A., Hertel, K.J. and Shi, Y. (2011) Complex and dynamic landscape of RNA polyadenylation revealed by PAS-Seq. RNA, 17, 761–772.

8. Bennett, D.A., Schneider, J.A., Buchman, A.S., de Leon, C.M., Bienias, J.L. and Wilson, R.S. (2005) The Rush Memory and Aging Project: study design and baseline characteristics of the study cohort. Neuroepidemiology, 25, 163–175.

9. Bennett, D.A., Schneider, J.A., Arvanitakis, Z. and Wilson, R.S. (2012) OVERVIEW AND FINDINGS FROM THE RELIGIOUS ORDERS STUDY. Curr Alzheimer Res, 9, 628.

10. Bennett, D.A., Schneider, J.A., Buchman, A.S., Barnes, L.L., Boyle, P.A. and Wilson, R.S. (2012) Overview and Findings from the Rush Memory and Aging Project. Curr Alzheimer Res, 9, 646.

11. Wang, Z., Jensen, M.A. and Zenklusen, J.C. (2016) A Practical Guide to The Cancer Genome Atlas (TCGA). Methods Mol Biol, 1418, 111–141.

12. Baxi, E.G., Thompson, T., Li, J., Kaye, J.A., Lim, R.G., Wu, J., Ramamoorthy, D., Lima, L., Vaibhav, V., Matlock, A., et al. (2022) Answer ALS, a large-scale resource for sporadic and familial ALS combining clinical and multi-omics data from induced pluripotent cell lines. Nat Neurosci, 25, 226–237.

13. Chen, M., Ji, G., Fu, H., Lin, Q., Ye, C., Ye, W., Su, Y. and Wu, X. (2019) A survey on identification and quantification of alternative polyadenylation sites from RNA-seq data. Brief Bioinform, 21, 1261–1276.

14. Lee, J.Y., Yeh, I., Park, J.Y. and Tian, B. (2007) PolyA_DB 2: mRNA polyadenylation sites in vertebrate genes. Nucleic Acids Res, 35.

15. Wu, X., Zhang, Y. and Li, Q.Q. (2016) PlantAPA: A Portal for Visualization and Analysis of Alternative Polyadenylation in Plants. Front Plant Sci, 7.

16. Gruber, A.J., Schmidt, R., Gruber, A.R., Martin, G., Ghosh, S., Belmadani, M., Keller, W. and Zavolan, M. (2016) A comprehensive analysis of 3’ end sequencing data sets reveals novel polyadenylation signals and the repressive role of heterogeneous ribonucleoprotein C on cleavage and polyadenylation. Genome Res, 26, 1145–1159.

17. Wang, R., Nambiar, R., Zheng, D. and Tian, B. (2018) PolyA_DB 3 catalogs cleavage and polyadenylation sites identified by deep sequencing in multiple genomes. Nucleic Acids Res, 46, D315–D319.

18. Herrmann, C.J., Schmidt, R., Kanitz, A., Artimo, P., Gruber, A.J. and Zavolan, M. (2020) PolyASite 2.0: a consolidated atlas of polyadenylation sites from 3’ end sequencing. Nucleic Acids Res, 48, D174–D179.

19. Wang, R. and Tian, B. (2020) APAlyzer: A bioinformatics package for analysis of alternative polyadenylation isoforms. Bioinformatics, 36, 3907–3909.

20. Grassi, E., Mariella, E., Lembo, A., Molineris, I. and Provero, P. (2016) Roar: Detecting alternative polyadenylation with standard mRNA sequencing libraries. BMC Bioinformatics, 17, 1–9.

21. Ha, K.C.H., Blencowe, B.J. and Morris, Q. (2018) QAPA: A new method for the systematic analysis of alternative polyadenylation from RNA-seq data. Genome Biol, 19, 1–18.

22. le Pera, L., Mazzapioda, M. and Tramontano, A. (2015) 3USS: a web server for detecting alternative 3’UTRs from RNA-seq experiments. Bioinformatics, 31, 1845–1847.

23. Huang, Z. and Teeling, E.C. (2017) ExUTR: A novel pipeline for large-scale prediction of 3’-UTR sequences from NGS data. BMC Genomics, 18, 1–11.

24. Birol, I., Raymond, A., Chiu, R., Nip, K.M., Jackman, S.D., Kreitzman, M., Docking, T.R., Ennis, C.A., Robertson, G.A. and Karsan, A. (2015) KLEAT: CLEAVAGE SITE ANALYSIS OF TRANSCRIPTOMES. Pac Symp Biocomput, 10.1142/9789814644730_0034.

25. Xia, Z., Donehower, L.A., Cooper, T.A., Neilson, J.R., Wheeler, D.A., Wagner, E.J. and Li, W. (2014) Dynamic analyses of alternative polyadenylation from RNA-seq reveal a 3’2-UTR landscape across seven tumour types. Nat Commun, 5.

26. Ye, C., Long, Y., Ji, G., Li, Q.Q. and Wu, X. (2018) APAtrap: Identification and quantification of alternative polyadenylation sites from RNA-seq data. Bioinformatics, 34, 1841–1849.

27. Arefeen, A., Liu, J., Xiao, X. and Jiang, T. (2018) TAPAS: tool for alternative polyadenylation site analysis. Bioinformatics, 34, 2521–2529.

28. Ji, Y., Zhou, Z., Liu, H. and Davuluri, R. v (2021) DNABERT: pre-trained Bidirectional Encoder Representations from Transformers model for DNA-language in genome. Bioinformatics, 10.1093/bioinformatics/btab083.

29. Dobin, A., Davis, C.A., Schlesinger, F., Drenkow, J., Zaleski, C., Jha, S., Batut, P., Chaisson, M. and Gingeras, T.R. (2013) STAR: ultrafast universal RNA-seq aligner. Bioinformatics, 29, 15.

30. Danecek, P., Bonfield, J.K., Liddle, J., Marshall, J., Ohan, V., Pollard, M.O., Whitwham, A., Keane, T., McCarthy, S.A., Davies, R.M., et al. (2021) Twelve years of SAMtools and BCFtools. Gigascience, 10.

31. Benjamini, Y., Drai, D., Elmer, G., Kafkafi, N. and Golani, I. (2001) Controlling the false discovery rate in behavior genetics research. Behavioural brain research, 125, 279–284.

32. Yalamanchili, H.K., Alcott, C.E., Ji, P., Wagner, E.J., Zoghbi, H.Y. and Liu, Z. (2020) PolyA-miner: Accurate assessment of differential alternative poly-adenylation from 3’Seq data using vector projections and non-negative matrix factorization. Nucleic Acids Res, 48, 1–12.

33. Ramírez, F., Bhardwaj, V., Arrigoni, L., Lam, K.C., Grüning, B.A., Villaveces, J., Habermann, B., Akhtar, A. and Manke, T. (2018) High-resolution TADs reveal DNA sequences underlying genome organization in flies. Nature Communications 2018 9:1, 9, 1–15.

34. Lopez-Delisle, L., Rabbani, L., Wolff, J., Bhardwaj, V., Backofen, R., Grüning, B., Ramírez, F. and Manke, T. (2021) pyGenomeTracks: reproducible plots for multivariate genomic datasets. Bioinformatics, 37, 422–423.

35. Hunter, J.D. (2007) Matplotlib: A 2D graphics environment. Comput Sci Eng, 9, 90–95.

36. de Maio, A., Yalamanchili, H.K., Adamski, C.J., Gennarino, V.A., Liu, Z., Qin, J., Jung, S.Y., Richman, R., Orr, H. and Zoghbi, H.Y. (2018) RBM17 Interacts with U2SURP and CHERP to Regulate Expression and Splicing of RNA-Processing Proteins. Cell Rep, 25, 726-736.e7.

37. Arora, A., Goering, R., Lo, H.Y.G., Lo, J., Moffatt, C. and Taliaferro, J.M. (2022) The Role of Alternative Polyadenylation in the Regulation of Subcellular RNA Localization. Front Genet, 12, 2791.

38. Zhang, L., Yan, F., Li, L., Fu, H., Song, D., Wu, D. and Wang, X. (2021) New focuses on roles of communications between endoplasmic reticulum and mitochondria in identification of biomarkers and targets. Clin Transl Med, 11.

39. Mak, H.Y., Ouyang, Q., Tumanov, S., Xu, J., Rong, P., Dong, F., Lam, S.M., Wang, X., Lukmantara, I., Du, X., et al. (2021) AGPAT2 interaction with CDP-diacylglycerol synthases promotes the flux of fatty acids through the CDP-diacylglycerol pathway. Nat Commun, 12.

40. Aypek, H., Krisp, C., Lu, S., Liu, S., Kylies, D., Kretz, O., Wu, G., Moritz, M., Amann, K., Benz, K., et al. (2022) Loss of the collagen IV modifier prolyl 3-hydroxylase 2 causes thin basement membrane nephropathy. J Clin Invest, 132.

41. Schulten, H.J., Al-Adwani, F., Saddeq, H.A.B., Alkhatabi, H., Alganmi, N., Karim, S., Hussein, D., Al-Ghamdi, K.B., Jamal, A., Al-Maghrabi, J., et al. (2022) Meta-analysis of whole-genome gene expression datasets assessing the effects of IDH1 and IDH2 mutations in isogenic disease models. Sci Rep, 12.

42. Fujiwara, T., Ye, S., Castro-Gomes, T., Winchell, C.G., Andrews, N.W., Voth, D.E., Varughese, K.I., Mackintosh, S.G., Feng, Y., Pavlos, N., et al. (2016) PLEKHM1/DEF8/RAB7 complex regulates lysosome positioning and bone homeostasis. JCI Insight, 1.

43. Yi, Y. and Ge, S. (2022) Targeting the histone H3 lysine 79 methyltransferase DOT1L in MLL-rearranged leukemias. J Hematol Oncol, 15.

44. Brumbaugh, J., di Stefano, B., Wang, X., Borkent, M., Forouzmand, E., Clowers, K.J., Ji, F., Schwarz, B.A., Kalocsay, M., Elledge, S.J., et al. (2018) Nudt21 Controls Cell Fate by Connecting Alternative Polyadenylation to Chromatin Signaling. Cell, 172, 106-120.e21.

45. Rüegsegger, U., Blank, D. and Keller, W. (1998) Human Pre-mRNA Cleavage Factor Im Is Related to Spliceosomal SR Proteins and Can Be Reconstituted In Vitro from Recombinant Subunits. Mol Cell, 1, 243–253.

46. Li, W., You, B., Hoque, M., Zheng, D., Luo, W., Ji, Z., Park, J.Y., Gunderson, S.I., Kalsotra, A., Manley, J.L., et al. (2015) Systematic Profiling of Poly(A)+ Transcripts Modulated by Core 3’ End Processing and Splicing Factors Reveals Regulatory Rules of Alternative Cleavage and Polyadenylation. PLoS Genet, 11, e1005166.

47. Martin, G., Gruber, A.R., Keller, W. and Zavolan, M. (2012) Genome-wide Analysis of Pre-mRNA 3’ End Processing Reveals a Decisive Role of Human Cleavage Factor I in the Regulation of 3’ UTR Length. Cell Rep, 1, 753–763.

48. Masamha, C.P., Xia, Z., Yang, J., Albrecht, T.R., Li, M., Shyu, A. bin, Li, W. and Wagner, E.J. (2014) CFIm25 links alternative polyadenylation to glioblastoma tumour suppression. Nature 2014 510:7505, 510, 412–416.

49. Weng, T., Huang, J., Wagner, E.J., Ko, J., Wu, M., Wareing, N.E., Xiang, Y., Chen, N.-Y., Ji, P., Molina, J.G., et al. (2020) Downregulation of CFIm25 amplifies dermal fibrosis through alternative polyadenylation. J Exp Med, 217.

50. Brown, K.M. and Gilmartin, G.M. (2003) A Mechanism for the Regulation of Pre-mRNA 3’ Processing by Human Cleavage Factor Im. Mol Cell, 12, 1467–1476.

51. Dong, Y., Fan, X., Wang, Z., Zhang, L. and Guo, S. (2020) Circ_HECW2 functions as a miR-30e-5p sponge to regulate LPS-induced endothelial-mesenchymal transition by mediating NEGR1 expression. Brain Res, 1748.

52. Krishnamoorthy, V., Khanna, R. and Parnaik, V.K. (2018) E3 ubiquitin ligase HECW2 mediates the proteasomal degradation of HP1 isoforms. Biochem Biophys Res Commun, 503, 2478–2484.

53. Krishnamoorthy, V., Khanna, R. and Parnaik, V.K. (2018) E3 ubiquitin ligase HECW2 targets PCNA and lamin B1. Biochim Biophys Acta Mol Cell Res, 1865, 1088–1104.

54. Stern, E.P., Guerra, S.G., Chinque, H., Acquaah, V., González-Serna, D., Ponticos, M., Martin, J., Ong, V.H., Khan, K., Nihtyanova, S.I., et al. (2020) Analysis of Anti-RNA Polymerase III Antibody-positive Systemic Sclerosis and Altered GPATCH2L and CTNND2 Expression in Scleroderma Renal Crisis. J Rheumatol, 47, 1668–1677.

55. Iyama, T. and Wilson, D.M. (2013) DNA repair mechanisms in dividing and non-dividing cells. DNA Repair (Amst), 12, 620–636.

56. Mangleburg, C.G., Wu, T., Yalamanchili, H.K., Guo, C., Hsieh, Y.C., Duong, D.M., Dammer, E.B., de Jager, P.L., Seyfried, N.T., Liu, Z., et al. (2020) Integrated analysis of the aging brain transcriptome and proteome in tauopathy. Mol Neurodegener, 15, 1–17.

57. r, de S.A., Lo, P., Em, M. and c, V. (2009) Global signatures of protein and mRNA expression levels. Mol Biosyst, 5, 1512–1526.

58. Alcott, C., Yalamanchili, H.K., Ji, P., van der Heijden, M., Saltzman, A., Leng, M., Bhatt, B., Hao, S., Wang, Q., Saliba, A., et al. (2019) Partial loss of CFIm25 causes aberrant alternative polyadenylation and learning deficits. 10.1101/735597.

59. ben Zablah, Y., Merovitch, N. and Jia, Z. (2020) The Role of ADF/Cofilin in Synaptic Physiology and Alzheimer’s Disease. Front Cell Dev Biol, 8.

60. Khan, R., Kulasiri, D. and Samarasinghe, S. (2021) Functional repertoire of protein kinases and phosphatases in synaptic plasticity and associated neurological disorders. Neural Regen Res, 16, 1150–1157.

61. Gns, H.S., Rajalekshmi, S.G. and Burri, R.R. (2022) Revelation of Pivotal Genes Pertinent to Alzheimer’s Pathogenesis: A Methodical Evaluation of 32 GEO Datasets. J Mol Neurosci, 72, 303–322.

62. Yoshino, Y., Yamazaki, K., Ozaki, Y., Sao, T., Yoshida, T., Mori, T., Mori, Y., Ochi, S., Iga, J.I. and Ueno, S.I. (2017) INPP5D mRNA Expression and Cognitive Decline in Japanese Alzheimer’s Disease Subjects. J Alzheimers Dis, 58, 687–694.

63. Liu, T., Zhu, B., Liu, Y., Zhang, X., Yin, J., Li, X., Jiang, L.L., Hodges, A.P., Rosenthal, S.B., Zhou, L., et al. (2020) Multi-omic comparison of Alzheimer’s variants in human ESC-derived microglia reveals convergence at APOE. J Exp Med, 217.

64. Tsai, A.P., Lin, P.B.C., Dong, C., Moutinho, M., Casali, B.T., Liu, Y., Lamb, B.T., Landreth, G.E., Oblak, A.L. and Nho, K. (2021) INPP5D expression is associated with risk for Alzheimer’s disease and induced by plaque-associated microglia. Neurobiol Dis, 153.

65. Kim, J.H. (2018) Genetics of Alzheimer’s Disease. Dement Neurocogn Disord, 17, 131.

66. Karch, C.M. and Goate, A.M. (2015) Alzheimer’s disease risk genes and mechanisms of disease pathogenesis. Biol Psychiatry, 77, 43–51.

67. Akhtar, A. and Sah, S.P. (2020) Insulin signaling pathway and related molecules: Role in neurodegeneration and Alzheimer’s disease. Neurochem Int, 135.

68. Kellar, D. and Craft, S. (2020) Brain insulin resistance in Alzheimer’s disease and related disorders: mechanisms and therapeutic approaches. Lancet Neurol, 19, 758–766.

69. Wu, H., Dunnett, S., Ho, Y.S. and Chang, R.C.C. (2019) The role of sleep deprivation and circadian rhythm disruption as risk factors of Alzheimer’s disease. Front Neuroendocrinol, 54.

70. Uddin, M.S., Tewari, D., Mamun, A. al, Kabir, M.T., Niaz, K., Wahed, M.I.I., Barreto, G.E. and Ashraf, G.M. (2020) Circadian and sleep dysfunction in Alzheimer’s disease. Ageing Res Rev, 60.

71. Choi, S.H., Bylykbashi, E., Chatila, Z.K., Lee, S.W., Pulli, B., Clemenson, G.D., Kim, E., Rompala, A., Oram, M.K., Asselin, C., et al. (2018) Combined adult neurogenesis and BDNF mimic exercise effects on cognition in an Alzheimer’s mouse model. Science, 361.

72. Amidfar, M., de Oliveira, J., Kucharska, E., Budni, J. and Kim, Y.K. (2020) The role of CREB and BDNF in neurobiology and treatment of Alzheimer’s disease. Life Sci, 257.

73. Zhang, H., Zhang, L., Zhou, D., Li, H. and Xu, Y. (2021) ErbB4 mediates amyloid β-induced neurotoxicity through JNK/tau pathway activation: Implications for Alzheimer’s disease. J Comp Neurol, 529, 3497–3512.

74. Ou, G.Y., Lin, W.W. and Zhao, W.J. (2021) Neuregulins in Neurodegenerative Diseases. Front Aging Neurosci, 13.

75. Lim, N.S., Swanson, C.R., Cherng, H.R., Unger, T.L., Xie, S.X., Weintraub, D., Marek, K., Stern, M.B., Siderowf, A., Trojanowski, J.Q., et al. (2016) Plasma EGF and cognitive decline in Parkinson’s disease and Alzheimer’s disease. Ann Clin Transl Neurol, 3, 346–355.

76. Petrelis, A.M., Stathopoulou, M.G., Kafyra, M., Murray, H., Masson, C., Lamont, J., Fitzgerald, P., Dedoussis, G., Yen, F.T. and Visvikis-Siest, S. (2022) VEGF-A-related genetic variants protect against Alzheimer’s disease. Aging, 14, 2524–2536.

