## Supplemental Figures for "PolyAMiner-Bulk: A Machine Learning Based Bioinformatics Algorithm to Infer and Decode Alternative Polyadenylation Dynamics from bulk RNA-seq data"

**SUPPLEMENTARY FIGURE S1**

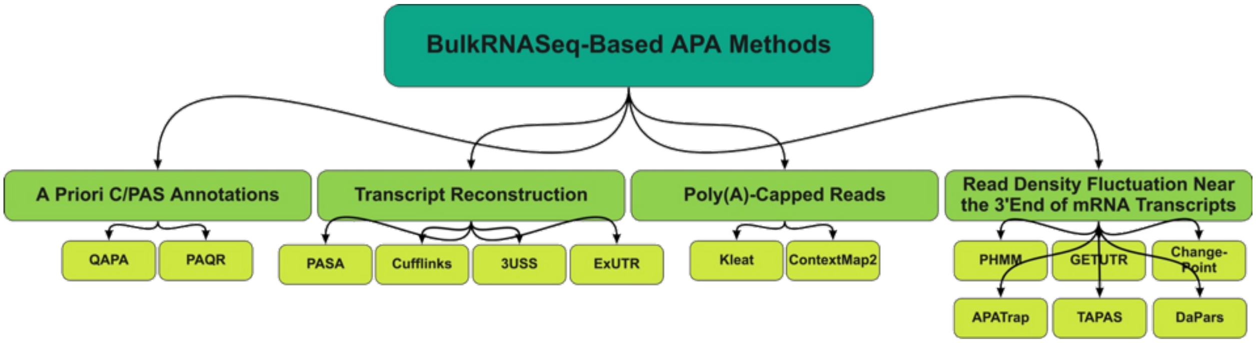

### SUPPLEMENTARY FIGURE S2

**DEF8 = Representative Differential APA Gene (DAG)**

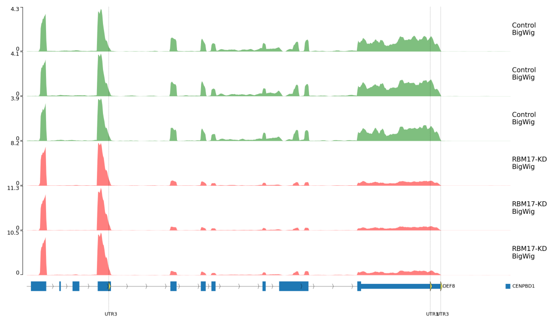


**Bulk RNA-Seq Read Density Visualization**

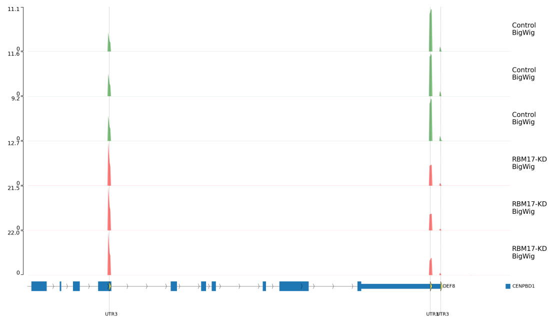


**Pseudo 3’-Seq Read Density Visualization**

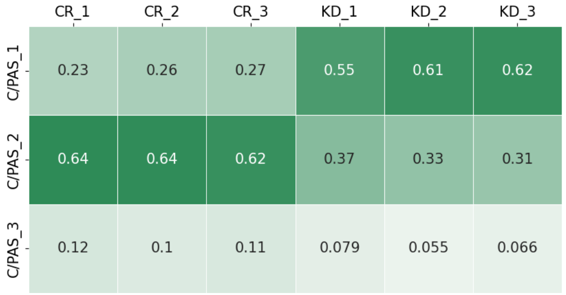


**Read Density Heatmap**

### SUPPLEMENTARY FIGURE S3

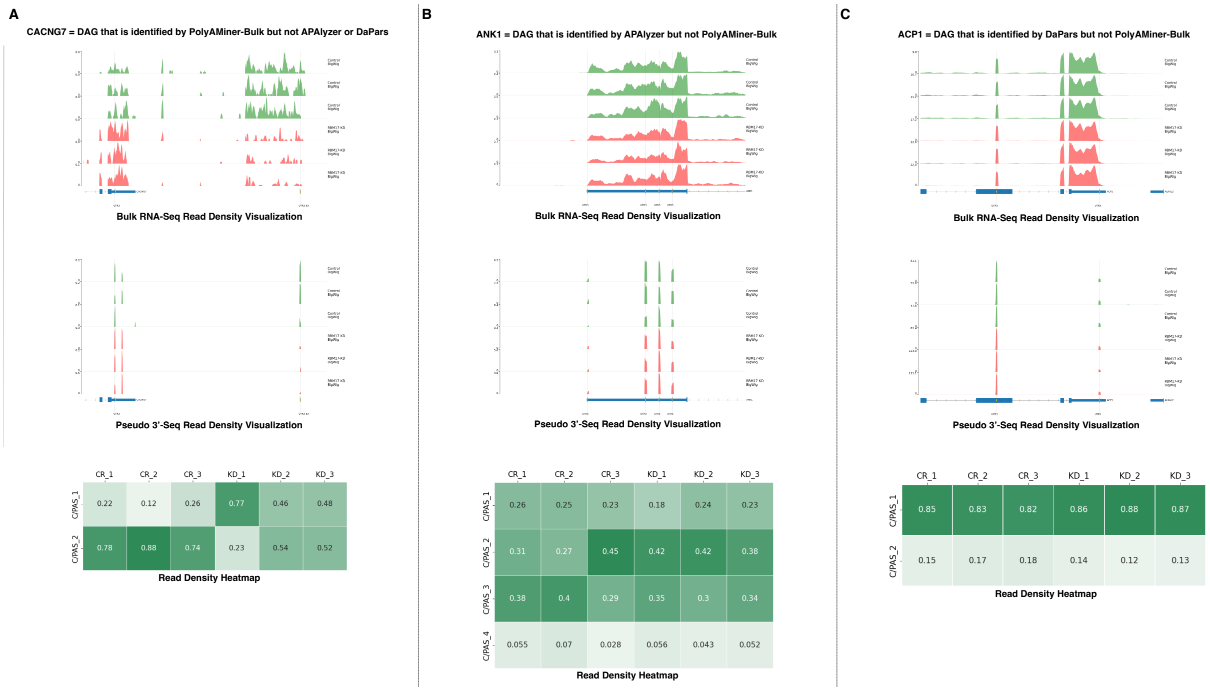

#### SUPPLEMENTARY FIGURE S4

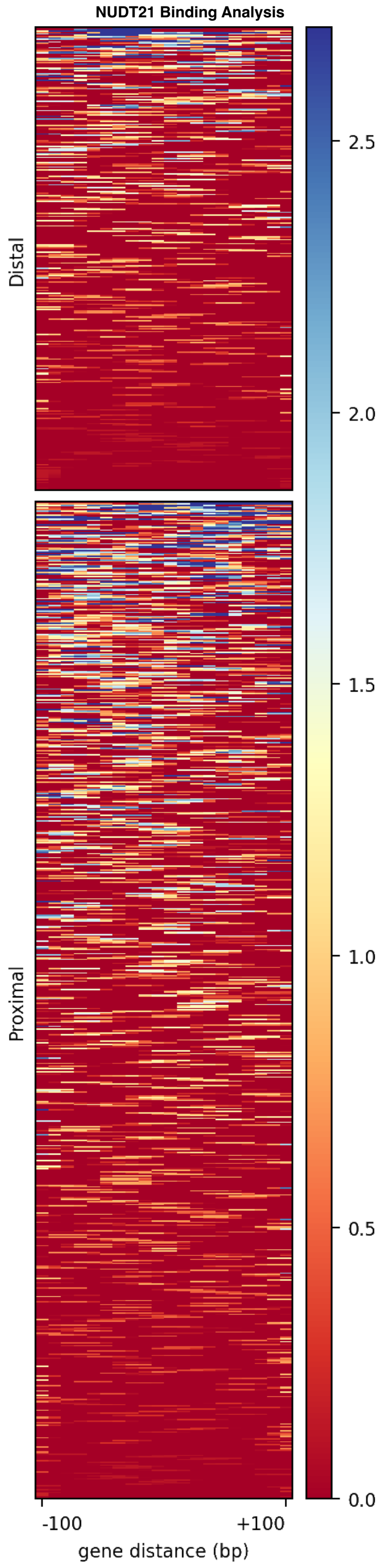

**Supplementary Table S1**

| Feature | PolyAMiner-Bulk | APAlyzer | APAtrap | DaPars | QAPA | Roar | TAPAS |
|---|---|---|---|---|---|---|---|
| *De novo* C/PAS identification | **YES** | NO | **YES** | **YES** | NO | NO | **YES** |
| Reference Database | **PolyASite & PolyA_DB** | **PolyA_DB only** | N/A | N/A | **PolyASite/ GENCODE** | **PolyA_DB & APASdb** | N/A |
| Deep Learning Model | **YES** | NO | NO | NO | NO | NO | NO |
| Intral-distal and intra-proximal APA quantification | **YES** | NO | NO | NO | NO | NO | NO |
| 3'UTR APA | **YES** | **YES** | **YES** | **YES** | **YES** | **YES** | **YES** |
| IPA | **YES** | **YES** | NO | NO | NO | NO | NO |
| Visualization (Volcano Plot) | **YES** | **YES** | NO | NO | NO | NO | NO |
| Visualization (IGV Read Density) | **YES** | NO | NO | NO | NO | NO | NO |
| Visualization (Heatmap) | **YES** | NO | NO | NO | NO | NO | NO |
| Reference | This study | (Wang, et al., 2020) | (Ye, et al., 2018) | (Xia, et al., 2014) | (Ha, et al., 2018) | (Grassi, et al., 2016) | (Arefeen, et al., 2018) |

**Program download site:**
PolyAMiner-Bulk: https://github.com/venkatajonnakuti/PolyAMiner-Bulk
APAlyzer: https://bioconductor.org/packages/release/bioc/html/APAlyzer.html
APAtrap: https://sourceforge.net/projects/apatrap/
DaPars: https://github.com/ZhengXia/dapars
QAPA: https://github.com/morrislab/qapa
ROAR: https://bioconductor.org/packages/release/bioc/html/roar.html
TAPAS: https://github.com/arefeen/TAPAS

**References:**
Wang, R., et al. APAlyzer: a bioinformatics package for analysis of alternative poladenylation isoforms. Bioinformatics 2020; 36(12):3907-3309
Arefeen, A., et al. TAPAS: tool for alternative polyadenylation site analysis. Bioinformatics 2018;34(15):2521-2529.
Grassi, E., et al. Roar: detecting alternative polyadenylation with standard mRNA sequencing libraries. BMC bioinformatics 2016;17(1):423.
Ha, K.C., Blencowe, B.J. and Morris, Q. QAPA: a new method for the systematic analysis of alternative polyadenylation from RNA-seq data. Genome biology 2018;19(1):45.
Xia, Z., et al. Dynamic analyses of alternative polyadenylation from RNA-seq reveal a 3′-UTR landscape across seven tumour types. Nature communications 2014;5(1):1-13.
Ye, C., et al. APAtrap: identification and quantification of alternative polyadenylation sites from RNA-seq data. Bioinformatics 2018;34(11):1841-1849.
